# Sparse Project VCF: efficient encoding of population genotype matrices

**DOI:** 10.1101/611954

**Authors:** Michael F. Lin, Xiaodong Bai, William J. Salerno, Jeffrey G. Reid

**Affiliations:** mlin.net LLC, San Jose, CA 95113, USA; Regeneron Genetics Center, Tarrytown, NY 10591, USA

## Abstract

**Summary:** Variant Call Format (VCF), the prevailing representation for germline genotypes in population sequencing, suffers rapid size growth as larger cohorts are sequenced and more rare variants are discovered. We present Sparse Project VCF (spVCF), an evolution of VCF with judicious entropy reduction and run-length encoding, delivering >10X size reduction for modern studies with practically minimal information loss. spVCF interoperates with VCF efficiently, including tabix-based random access.

**Availability and Implementation:** Freely available at github.com/mlin/spVCF

## 1 Introduction

Variant Call Format (VCF) is the prevailing representation for germline variants discovered by high-throughput sequencing (Danecek et al., 2011). In addition to capturing variants sequenced in one study participant, VCF can represent the genotypes for many participants at all discovered variant loci. This “Project VCF” (pVCF) form is a 2-D matrix with loci down the rows and participants across the columns, filled in with each called genotype and associated quality-control (QC) measures, such as read depths, read strand ratios, and genotype likelihoods.

As the number of study participants *N* grows (columns), more variant loci are also discovered (rows), leading to super-linear growth of the pVCF genotype matrix. And, because cohort sequencing discovers mostly rare variants, this matrix consists largely of reference-identical genotypes and their high-entropy QC measures. In recent experiments with human whole-exome sequencing (WES), doubling N from 25 000 to 50000 also increased the pVCF locus count by 43%, and 96% of all loci had nonreference allele frequency below 0.1% (Lin *et al*., 2018). Empirically, vcf.gz file sizes in WES and whole-genome sequencing (WGS) are growing roughly with *N*^1.5^ in the largest studies as of this writing (*N* ≈ 500 000 WES). Unchecked, we project *N* = 1000 000 WGS will yield petabytes of *compressed* pVCF.

## 2 Approach

We sought an incremental path to ameliorate the QC entropy and size growth problems in existing pVCF-based pipelines, which may be reluctant to adopt fundamentally different formats or data models addressing these challenges (Layer *et al*., 2015; Li, 2015; Zheng *et al*., 2017; LeFaive, 2017; Danek and Deorowicz, 2018; Klarqvist, 2018; Deorowicz and Danek, 2019) due to upstream & downstream tool dependencies. To this end, we developed an evolution of VCF, Sparse Project VCF (spVCF), which begins with the same data model and text format, and adds three simple ideas (**Fig. 1**):

1. *Squeezing: judiciously reducing QC entropy*. In any cell with zero reads supporting a variant (typically Allele Depth `AD` = *d,* 0 for any *d,* depending on the upstream caller) and corresponding reference-identical or non-called genotype, we discard all fields except the genotype `GT` and the read depth `DP`, which we also round down to a power of two (0, 1, 2, 4, 8, 16,…; configurable). This convention, inspired by similar techniques for base quality scores (Fritz *et al*., 2011; Illumina, 2014; Jun *et al*., 2015; Bonfield *et al*., 2018), aims to preserve nearly all *useful* information, removing uninformative fluctuations in QC measures. It leaves unmodified all cells with evidence for a variant, whether or not a variant genotype is actually called; in other cells, it maintains a binned lower bound on the reference depth.
2. *Succinct, lossless encoding for runs of reference-identical cells*. First we replace the contents of a reference-identical (or non-called) cell with a quotation mark “ if it’s identical to the cell above it, compressing runs down the column for each sample. Then we run-length encode these quotation marks across the rows, so for example a stretch of 42 marks across a row is written `<tab>“42` instead of repeating `<tab>”` forty-two times. (This second step has negligible effect on zipped file size, but allows efficient processing of the encoded form without expansion.) The QC squeezing synergistically enables the run-encoding, by converting small fluctuations in closely-spaced cells into identical runs down each sample column.
3. *Checkpointing to facilitate random access* by genome range (row) within a spVCF file. While all *variant* genotype cells are readily accessible from a given spVCF row, fully decoding the reference-identical and noncalled cells would require information from an unpredictable number of prior rows. To expedite random access, the spVCF encoder periodically skips run-encoding a row, instead emitting a row identical to the squeezed pVCF. Subsequent run-encoded rows can be decoded by looking back no farther than this *checkpoint* row. Every run-encoded row has an additional informational field with the position of the previous checkpoint. Genome range access proceeds by locating the first desired row, following its pointer back to a checkpoint, and reversing the run-encoding from the checkpoint through the desired row(s).

**Fig. 1.**
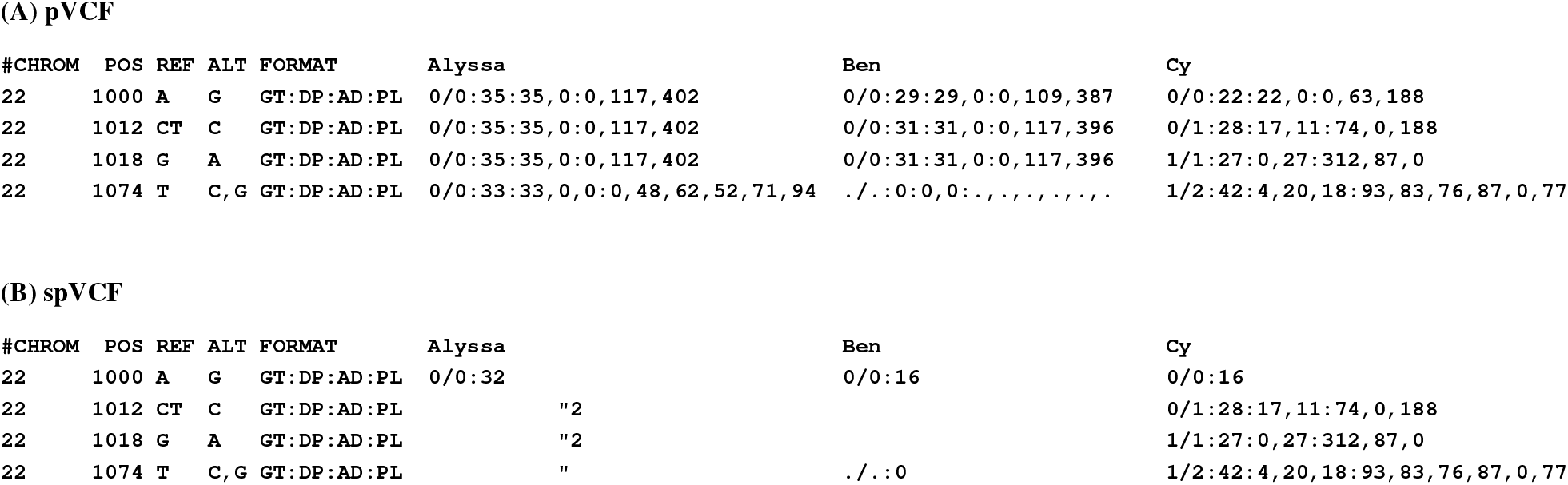
spVCF encoding example. (A) Illustrative pVCF of four variant loci in three sequenced study participants, with matrix entries encoding called genotypes and several numeric QC measures. Some required VCF fields are omitted for brevity. (B) spVCF encoding of the same example. QC values for reference-identical and non-called cells are reduced to a power-of-two lower bound on read depth DP. Runs of identical entries down columns are abbreviated using quotation marks, then runs of these marks across rows are length-encoded. Cy’s entries are shown column-aligned for clarity; the encoded text matrix is ragged.

## 3 Reference implementation

Our Unix command-line tool spvcf provides efficient transcoding between pVCF and spVCF, typically arranged in a shell pipeline to `gunzip` the input and `bgzip` the output. Different invocations of the tool can cause it to (i) squeeze and run-encode pVCF to spVCF, (ii) run-encode pVCF losslessly without squeezing, (iii) squeeze pVCF without run-encoding (producing valid pVCF that is typically much smaller, albeit not as small as spVCF), or (iv) decode spVCF back to pVCF.

If a spVCF file is compressed using bgzip, then tabix can create a random-access index for it (Li, 2011), as the encoding does not affect the necessary locus-level VCF fields. A subcommand of `spvcf` used instead of `tabix` can then access rows by genome position, consulting the checkpoints to formulate a spVCF “slice” that can be decoded standalone. The encoder checkpoints at a regular, configurable period and at the start of each chromosome; more-strategic checkpointing might improve compression slightly in the future.

The Apache-licensed code, compiled Linux executable, and detailed format documentation are available from: github.com/mlin/spVCF

## 4 Compression in DiscovEHR and UK Biobank

We tested spVCF on two sizeable WES studies using different upstream variant-calling pipelines.

First, using *N* = 50 000 WES from the DiscovEHR study (Dewey *et al*., 2016), we reduced a GATK-based pVCF file with 620 782 chromosome 2 variant loci from 79GiB `vcf.gz` to a 5.2GiB `spvcf.gz` file, 15X size reduction. Most of this reduction (6.9X) was achieved by the QC squeezing, while the run-encoding contributed an additional 2.2X. Experiments with nested subsets of these *N* = 50 000 WES indicate `spvcf.gz` file sizes growing roughly with *N*^1.1^, compared to *N*^1.5^ for the original `vcf.gz`. (All `.gz` files were made by bgzip on default compression level.) VCF’s binary equivalent, BCF, reduces this example by 1.2X losslessly and exhibits the same *N*^1.5^ scaling.

Second, with N = 302 342 WES from UK Biobank (Bycroft *et al*., 2018; Van Hout *et al*., 2019), spVCF reduced `vcf.gz` files for 252 610 loci in ten representative chromosome 2 segments from 110GiB to 7.7GiB. This 14X combined ratio is similar to that achieved for DiscovEHR; decomposed however, QC squeezing was relatively less impactful (4.2X) and run-encoding relatively moreso (3.4X). On the one hand, the UK Biobank pVCF files were produced using a different upstream pipeline (“SPB”) that already omitted genotype likelihoods for most reference-identical cells, leaving less to be squeezed out compared to DiscovEHR. On the other hand, the effectiveness of the run-encoding increased with the 3.3X-higher density of variant loci in the larger cohort, a trend expected to further amplify with larger *N*.

In these tests, `spvcf` squeezed and run-encoded the uncompressed pVCF at roughly the same rate that `bgzip` would compress the same input (each on a single x86-64 thread; both tools also have multithread modes). The decoder, with inputs and outputs both much smaller than the original pVCF, is several times faster. This makes it feasible to store compact spVCF files and decode them to pVCF only for transient use with tools expecting that format. The smaller squeezed pVCF also tends to speed up such tools.

## 5 Discussion

spVCF is a practical “next step” for storage and transfer in ongoing cohort sequencing projects, owing to its identical data model and performant transcoding with pVCF, providing interoperability with existing pVCF-based tools. Upstream, joint-calling tools such as such as GATK GenotypeGVCFs (DePristo *et al*., 2011; Poplin *et al*., 2018) and GLnexus (Lin *et al*., 2018) can “pipe” generated pVCF data into `spvcf` for now, and perhaps evolve to generate spVCF, or at least squeezed pVCF, internally. Downstream, existing population analysis tools can stream decoded pVCF from `spvcf`, with the future possibility of specializing algorithms to use the run-encoded form without expansion.

spVCF’s far-reduced size growth – though still slightly super-linear in *N*, owing to multiallelic loci and residual depth fluctuations – clears the way to scale up the VCF data model to *N* = 1 000 000 WGS studies in the near future. Meanwhile, many investigators – motivated by continuing advances in linked- and long-read sequencing – are developing haplotype-centric paradigms which might eventually replace VCF.

## Acknowledgements

We thank the Global Alliance for Genomics and Health, Large-Scale Genomics Work Stream for motivating discussions and early feedback on this work; particularly Albert Smith, Yossi Farjoun, Louis Bergelson, Chris Vittal, Cotton Seed, Cristina Gonzalez, Petr Danecek, Marcus Klarqvist, Rishi Nag, Richard Durbin, Thomas Keane, and Ewan Birney.

## Manuscript revision history

2020-08-25: New *N*=300K UKB results, updated bgzip speed comparison, clarify approach and discussion, add GTShark citation, correct SeqArray citation (with thanks to Stephanie Gogarten) 2019-04-17: First bioRxiv version

